# Laser facilitated epicutaneous peptide immunization using dry patch technology

**DOI:** 10.1101/2021.04.27.441719

**Authors:** Sandra Scheiblhofer, Stephan Drothler, Werner Braun, Reinhard Braun, Maximilian Boesch, Richard Weiss

## Abstract

The skin has been intensely investigated as a target tissue for immunization because it is populated by multiple types of antigen presenting cells. Directly addressing dendritic cells or Langerhans cells *in vivo* represents an attractive strategy for inducing T cell responses in cancer immunotherapy. We and others have studied fractional laser ablation as a novel method combining efficient delivery of macromolecules to the skin with an inherent adjuvant effect of laser illumination. In this proof of concept study, we demonstrate the feasibility of peptide delivery to the skin using the P.L.E.A.S.E. professional Erb:YAG fractional infrared laser together with EPIMMUN patches. In an ovalbumin mouse model we demonstrate that a dry patch formulation of SIINFEKL peptide in combination with CpG-ODN1826, but not imiquimod or polyI:C, induces potent cytotoxic T cell responses, which can be further boosted by co-delivery of the pan-helper T cell epitope PADRE.

## Introduction

It has been previously demonstrated that macromolecules can be efficiently delivered to the upper layers of the skin via laser-generated micropores [1, 2]. Moreover, illumination with laser light using specific settings has been shown to significantly potentiate immune responses. This inherent adjuvant effect of laser treatment has been attributed to activation of dendritic cells at the irradiation site via release of specific “danger” signaling molecules from epithelial cells and an increase in dendritic cell motility [3, 4]. Whereas most of the studies employing laser beams for pre-treatment of vaccination sites merely investigated antibody responses and induction of T helper subsets, immunization of mice by laser-facilitated delivery of vaccinia virus encoding ovalbumin led to significantly higher numbers of IFN-γ secreting CD8 T cells compared to intradermally injected animals [5]. Furthermore, enhanced protection of pre-immunized mice against challenge with influenza virus was demonstrated [4]. However, in this case, laser treatment did not serve for vaccine delivery purpose, but was solely employed to exert adjuvant potency in combination with intradermal injections.

For decades, *ex vivo* generated dendritic cells of different subtypes, which had been isolated from patients, expanded, loaded with tumor cell lysates or antigens, and matured, have been used for cellular cancer immunotherapy [6]. More recently, *in vivo* delivery of antigens or adjuvants to DCs by targeting specific receptors has come into focus for tumor treatment. In this context, the richness of the skin in DCs renders this tissue an ideal target for both prophylactic and therapeutic vaccination. Previous studies investigating laser-facilitated skin vaccination in cancer models used laser illumination only as adjuvant before injection of *ex vivo* loaded DCs [7], or employed vaccibodies targeting whole antigen to specific DC subsets [2].

In contrast, in our preliminary study, we delivered OVA peptide 257-264 (SIINFEKL) to the skin via laser-generated micropores. The peptide was administered as a dry powder incorporated into a patch recently developed by Pantec Biosolutions AG and combined with TLR agonists CpG-ODN 1826, imiquimod, and poly:IC, which have been intensely studied as adjuvants for cancer therapy. Furthermore, we tested whether addition of the universal T helper epitope PADRE would increase the potential of the peptide vaccine to induce specific CD8+ T cells with *in vivo* cytotoxic activity.

## Materials and methods

### Reagents

PADRE peptide (AKFVAAWTLKAAA) and the immunodominant class I peptide of C57BL/6 mice (SIINFEKL) were synthesized by Bachem AG (Bubendorf, Switzerland). TLR agonists CpG-ODN 1826, imiquimod, and polyI:C HMW were from Invivogen (Toulouse, France). SIINFEKL-specific pentamers were from Proimmune (Oxford, UK).

### EPIMMUN patches

Dry vaccine patches (EPIMMUN patches) containing 100µg SIINFEKL or a mixture of SIINFEKL and 50µg of PADRE were manufactured by Pantec Biosolutions AG using a proprietary printing-like technology. EPIMMUN patches are occlusive and correspond to transdermal therapeutic systems (TTS) as specified in the European Pharmacopoeia.

### Mice and their housing

Wild-type female C57BL/6 mice aged between six and eight weeks were purchased from Janvier, France and maintained at the animal facility at the University of Salzburg under SPF (specific pathogen-free) conditions according to FELASA guidelines. Mice were used at 8-10 weeks of age for vaccination. All animal experiments were conducted in compliance with EU Directive 2010/63/EU and have been approved by the Austrian Ministry of Education, Science and Research, under permit number GZ 2020-0.282.314.

### Laser-assisted epicutaneous immunization

One day before laser microporation, the dorsal or ventral skin of the mice was depilated as previously described [8]. Epicutaneous immunization was performed with a P.L.E.A.S.E. ^®^ Professional laser system (Pantec Biosolutions AG) using the following parameters: 0.7W, three pulses, total fluence of 8.3J/cm^2^, 9% pore density, 1cm diameter as previously described [8].

50µg (experiment 1) or 20µg (experiment 2) of adjuvant were applied to the laserporation site in endotoxin-free PBS in a total volume of 40µL. Control groups without adjuvant received PBS alone. In case of imiquimod, due to lower solubility, a larger volume of up to 83µL was applied by repeating the procedure after the first 40µL had been taken up by the micropores. Mice were kept on a heating pad until the solution had been completely taken up via the micropores. The treated area was then covered with an EPIMMUN dry patch containing 100µg OVA peptide SIINFEKL or a mixture of SIINFEKL and 50µg of the pan DR T helper epitope PADRE. Patches were removed 24 hours later.

Mice were immunized twice at 14-(experiment 1) or 21-day intervals (experiment 2) and blood samples were taken 7 days after the first immunization and at the time-point of sacrifice (7 days after the 2^nd^ immunization).

### Immunological assays

For in *vitro* detection of SIINFEKL-specific CD8+ T cells, blood samples (50µL) from both time-points were stained with SIINFEKL pentamers (Proimmune, Oxford, UK) and analysed by flow cytometry according to the manufacturer’s instructions. At the endpoint, spleen and lymph node cells were prepared as previously described [1] and stained with pentamers following the manufacturer’s instructions.

*In vivo* CTL activity was assessed as previously described [9]. Briefly, splenocytes from naïve syngeneic C57BL/6 mice were prepared one day before sacrifice. Half of the cells were stained with fluorescent cell proliferation dye eFluor 450, the other with eFluor 670 (both from eBioscience) according to the manufacturer’s instructions. The eFluor 450 stained cells were subsequently pulsed with SIINFEKL peptide, whereas the other cells were left untreated. The two suspensions were mixed at equal ratios and adjusted to 3×10^7^ cells per mL. 100µL thereof were injected into the tail vein of immunized and control mice. 16 hours later, the recipient mice were sacrificed and their spleen and lymph node cells harvested. By flow cytometric analysis the percentage of specific lysis was determined as the ratio of peptide-pulsed (eFluor 450 stained) to un-pulsed (eFluor 670 stained) cells.

Additionally, cells were cultured in the presence of peptide overnight to determine the number of IFN-γ producing T cells by ELISPOT [9].

## Results

Mice were immunized epicutaneously via laser generated micropores using skin patches containing a dry formulation of the major class I epitope of OVA, SIINFEKL. The effect of different TLR agonists to boost cytotoxic T lymphocyte (CTL) responses against SIINFEKL was assessed.

As shown in Fig.1A, application of TLR9 agonist CpG-ODN 1826 resulted in a temporary decrease in body weight between day 7 and 14, but animals recovered after that. 14 days after the first immunization on the back, mice showed hyperpigmentation in the shaved and depilated area, which is due to a depilation-induced regenerative response of melanocyte stem cells [10]. Sites of laser-microporation showed delayed hair re-growth in some animals. Interestingly, CpG-ODN 1826 (and to a lesser extent polyI:C) more strongly inhibited hair re-growth and blocked hyperpigmentation at the site of immunization (Fig. 1B).

**Fig. 1.**
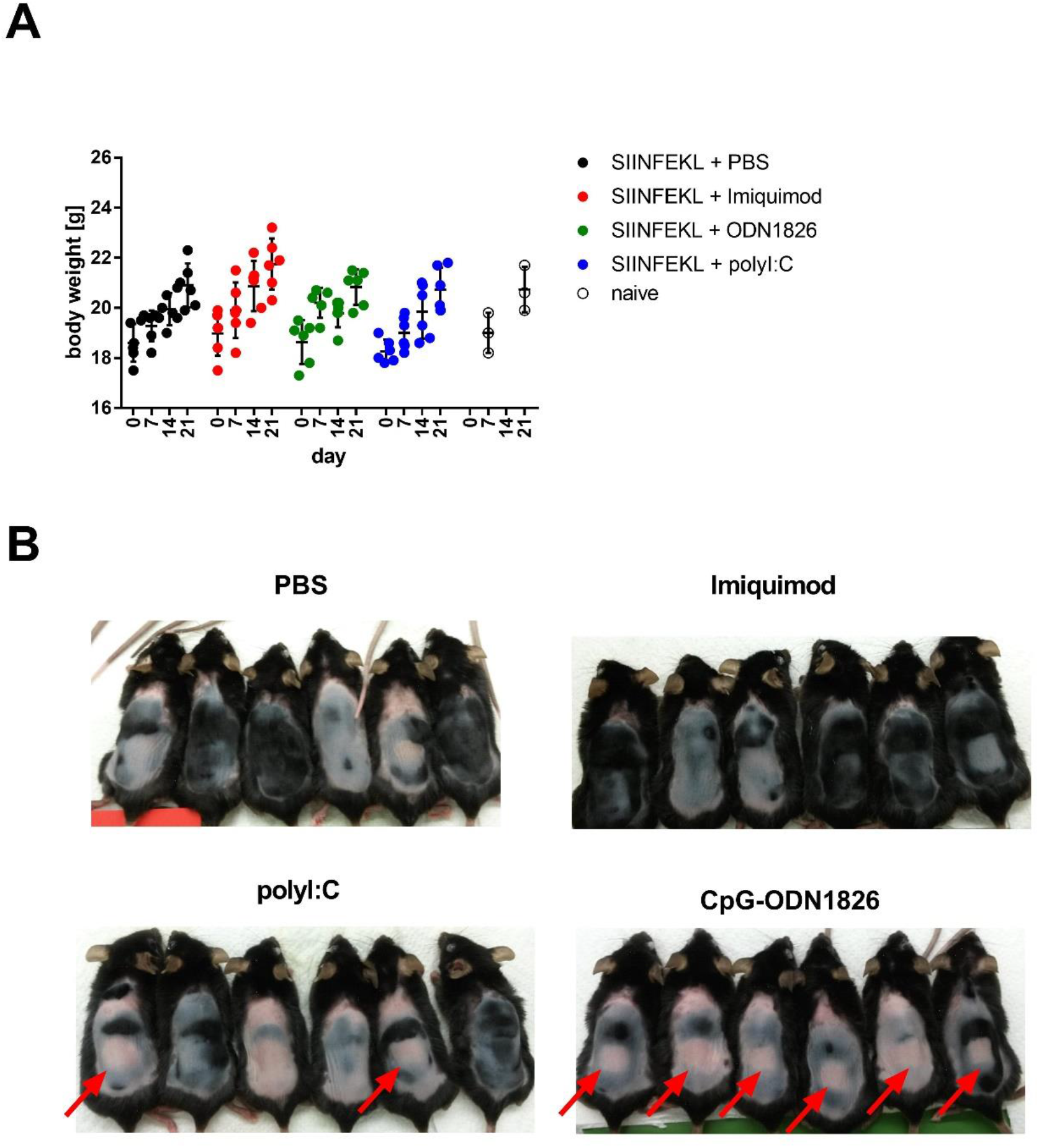
Effects of TLR-agonists on body weight (A) and skin pigmentation (B). Mice (n=6) were immunized on days 0 and 14 with 100µg SIINFEKL (dry patch) plus 50µg of different TLR-agonists or vehicle control (PBS). Naïve animals (n=3) served as control. 14 days after the first immunization, mice showed depilation induced hyperpigmentation. Red arrows indicate the square laser sites that displayed delayed hair re-growth and lack of hyperpigmentation.

Induction of CD8+ T cell responses was assessed by measuring the percentage of SIINFEKL-specific CD8+ T cells using class I pentamers specific for T cell receptor bound SIINFEKL in blood, skin draining lymph nodes (SDLN), and spleen (Fig. 2A). Furthermore, the number of IFN-γ producing splenocytes after stimulation with SIINFEKL was measured by ELISPOT (Fig. 2B). Finally, specific killing (lysis) of SIINFEKL-loaded target cells was evaluated using an *in vivo* CTL assay (Fig. 2C). As shown in Fig.2, in the absence of adjuvant (PBS control), no SIINFEKL-specific CD8+ T cells were found in the blood 7 days after the first and after the second immunization. Of the tested TLR agonists CpG-ODN 1826 (TLR9), imiquimod (TLR7/8), and polyI:C (TLR3), only CpG-ODN 1826 significantly boosted the number of SIINFEKL-specific CD8+ T cells circulating in blood after the first and second immunization. This also corresponded with significantly increased numbers of SIINFEKL-specific CD8+ T cells residing in the secondary lymphatic organs.

**Fig. 2.**
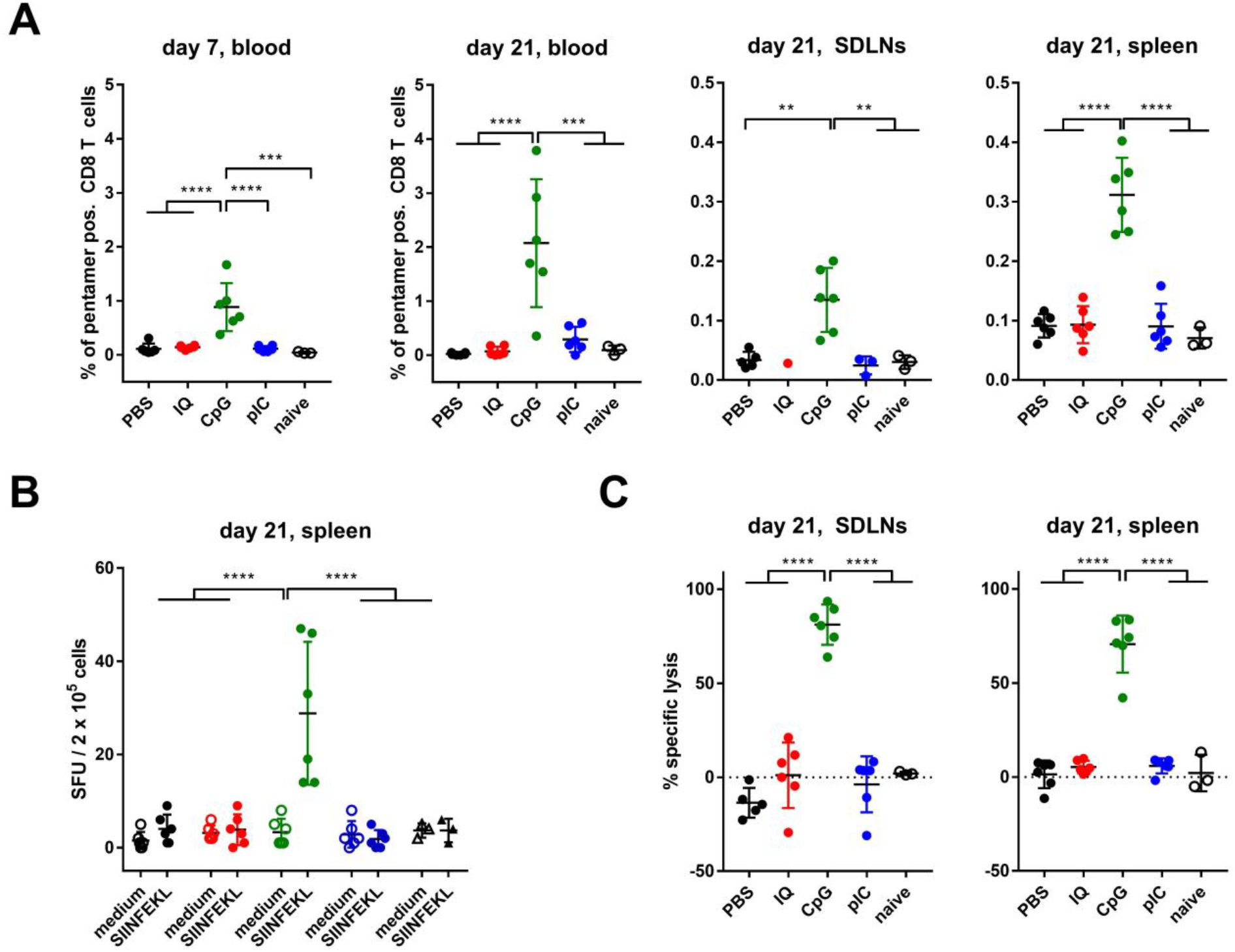
Effects of TLR-agonists on the immunogenicity of SIINFEKL-EPIMMUN patches. Mice (n=6) were immunized on days 0 and 14 with 100µg SIINFEKL (dry patch) plus 50µg of different TLR-agonists or vehicle control (PBS). A) The percentage of SIINFEKL-pentamer positive CD8+ T cells was determined in blood, skin draining lymph nodes, and spleen at the indicated days. B) Splenocytes from the same animals were analyzed by ELISPOT for the number of IFN-γ producing cells after 24h of in vitro re-stimulation with medium or SIINFEKL peptide. C) Cytotoxic T cells were analyzed using an *in vivo* functional CTL assay and specific lysis was determined by quantitating the number of SIINFEKL pulsed target cells (eFluor 450 label) vs. unpulsed target cells (eFluor 670 label) in skin draining lymph nodes and spleen. IQ = imiquimod, CpG = CpG-ODN1826, pIC = poly(I:C). Data are shown as means±SD and individual data points. Statistical significance was calculated by one-way (A and C) or RM two-way ANOVA (B) followed by Tukey’s post hoc test. ** P<0.01, *** P<0.001, **** P<0.0001. Missing data points in panel A, SDLNs are due to lack of sufficient quantities of cells.

However, the percentage of pentamer-positive resident CD8+ T cells was lower compared to the circulating CD8+ T cells, indicating that 7 days after the 2^nd^ immunization mainly circulating effector T cells rather than resident central memory T cells were observed. Secretion of IFN-γ from CD8+ T cells is an important proxy for their cytotoxic function. As shown by ELISPOT assay (Fig. 2B), only after stimulation with SIINFEKL (solid symbols), but not after stimulation with medium (open symbols), IFN-γ secreting cells were found in splenocyte cultures of mice immunized with SIINFEKL+CpG-ODN 1826, indicating the presence of functionally active, SIINFEKL-specific CTLs. This was confirmed by an *in vivo* CTL assay, where SIINFEKL-loaded splenocytes (target cells), or unloaded splenocytes (control cells) were injected into immunized mice, and the percentage of specifically killed target cells was assessed. As shown in Fig. 2C, mice immunized with SIINFEKL+CpG-ODN 1826 were able to kill SIINFEKL-loaded target cells with a very high efficacy.

At the time-point of sacrifice, mice immunized with CpG showed greatly enlarged lymph nodes and splenomegaly (data not shown), a known transient side effect of the adjuvant [11].

Our data has shown that in combination with a suitable adjuvant, a class I peptide alone applied via laser microporated skin is sufficient to induce a functional CTL response. However, it is known that CD4+ T cell help is needed for efficient priming [12] and generation of effector memory [13] CD8+ T cell responses. Consequently, it has been shown that addition of the pan-DR epitope PADRE can provide the necessary CD4+ T cell help to boost the efficacy of a therapeutic HPV-peptide vaccine [14]. This helper epitope can bind the most common HLA-DR molecules and also I-Ab molecules of C57BL/6 mice with high affinity and act as a powerful immunogen [15].

In consideration of this, we tested whether incorporation of PADRE into a dry patch formulation of SIINFEKL would boost CTL responses. As we have previously shown that 100µg SIINFEKL + 50µg CpG-ODN 1826 induced >90% CTL responses, we employed a lower concentration (20µg) of the adjuvant in this experiment to be able to discern a possible immune potentiating function of PADRE.

Immunization with 20µg CpG-ODN 1826 had no effect on hair regrowth or hyperpigmentation as seen with the 50µg dose (data not shown). The reduction of side effects was also evident in the absence of splenomegaly at the end of the experiment, although SDLNs were still larger compared to sham treated mice (data not shown). This confirms that the CpG-associated side effects are dose dependent. Levels of SIINFEKL-specific CD8+ T cells in blood were very heterogeneous, with individual mice displaying extremely high numbers (>20%) 7 days after the 2^nd^ immunization (Fig. 3A). ELISPOT assay and *in vivo* CTL clearly indicated that i) as expected, 20µg of CpG-ODN 1826 induced a weaker response compared to 50µg in the previous experiments; ii) co-immunization with PADRE alone was not sufficient to induce CTL responses; iii) co-immunization of PADRE together with 20µg CpG significantly boosted the CTL response to similar or higher levels as seen with 50µg CpG alone. As shown in Fig.3B, very high numbers of IFN-γ secreting cells were found in SDLNs and spleens of mice immunized with the SIINFEKL/PADRE dry patch together with 20µg CpG. These numbers were more than 5-fold higher (spleen) compared to immunization with SIINFEKL + CpG alone. This is in agreement with the *in vivo* CTL data that displayed an average killing capacity of ∼65% in the SIINFEKL+CpG group that was boosted to >90% by the addition of PADRE (Fig. 3C).

**Fig. 3.**
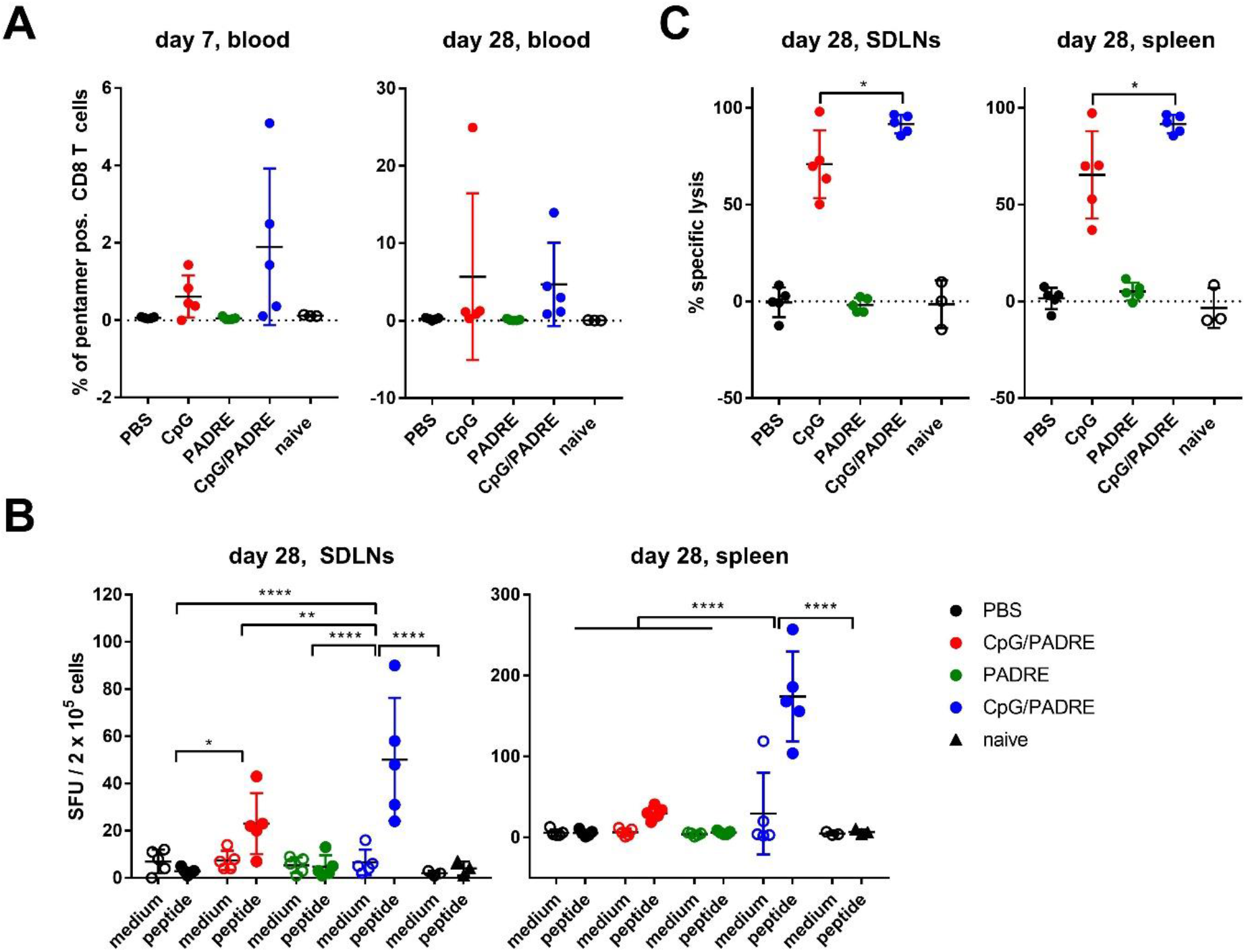
Effects of PADRE on SIINFEKL-EPIMMUN patches. Mice (n=5) were immunized on days 0 and 21 with 100µg SIINFEKL with or without 50µg PADRE (dry patch) plus 20µg of CpG-ODN1826 (CpG) or vehicle control (PBS). A) The percentage of SIINFEKL-pentamer positive CD8+ T cells was determined in blood at the indicated days. B) Splenocytes and cells from skin draining lymph nodes (SDLN) from the same animals were analyzed by ELISPOT for the number of IFN-γ producing cells after 24h of *in vitro* re-stimulation with medium or SIINFEKL peptide. C) Cytotoxic T cells were analyzed using an *in vivo* functional CTL assay and specific lysis was determined by quantitating the number of SIINFEKL pulsed target cells (eFluor 450 label) vs. unpulsed target cells (eFluor 670 label) in skin draining lymph nodes and spleen. Data are shown as means±SD and individual data points. Statistical significance was calculated by one-way (A and C) or RM two-way ANOVA (B) followed by Tukey’s post hoc test. * P<0.05, ** P<0.01, *** P<0.001, **** P<0.0001. For clarity, in panel C only statistical differences between the two CpG immunized groups are shown.

## Discussion and Conclusions

Dry formulations are advantageous for vaccines as their stability is greatly increased, allowing ambient transport and storage. Laser microporation is an ideal delivery method for dry powder vaccines because after treatment interstitial fluid enters the micropores and dissolves the powder vaccine from the applied patch, finally leading to uptake of the vaccine into the tissue [5]. Moreover, skin immunization methods frequently go along with localized side effects such as erythema, induration, edema or pruritus at the application site, especially when adjuvants are used. Therefore, novel delivery technologies and vaccine formulations, which minimize local side effects while maintaining vaccine immunogenicity and adjuvant potency, are needed.

Combining generation of skin micropores via laser irradiation with dry powder formulations has been shown to meet these requirements, as local adverse reactions were significantly reduced compared to intradermal injections [5].

In this proof of concept study, we could demonstrate that dry patch formulations of class I peptides applied to laser pre-treated skin can induce potent CTL responses that result in specific killing of >90% of injected target cells, when administered with CpG-ODN as adjuvant. This is in line with findings from Cheng et al., who showed that topical application of CpG is more potent in inducing CD8 responses than s.c. or i.m injections [16]. Although other studies showed that imiquimod [17, 18] or polyI:C [19] can boost SIINFEKL specific T cell responses after transcutaneous immunization, these TLR agonists failed to induce CTL responses in our study at the tested doses. This corroborates previous findings from our group, which showed that CpG, but not imiquimod or polyI:C suppress TH2 responses after laser-facilitated skin vaccination [1].

The receptor for CpG, TLR9, is differentially expressed in mice and men; i.e., whereas in mice TLR9 is expressed by all subsets of DCs, in humans it is restricted to plasmacytoid DCs, at least under steady state conditions [20]. However, there is clinical evidence that in tumor patients CpG can also drive activation of myeloid DCs and mature monocytes into DCs [21, 22], indicating that under inflammatory conditions, CpG may also target other types of DCs. Therefore, CpG motifs are still considered a promising adjuvant in humans, especially in the context of multi-modal therapy together with checkpoint blockers [23, 24].

In accordance with previous studies, we could show that the addition of a pan T helper epitope (PADRE) can significantly boost CTL responses when applied together with CpG-ODN 1826 [14, 25]. As this peptide binds to most HLA-DR alleles, it is a reasonable and feasible approach to incorporate this stimulant into personalized cancer vaccines.

Taken together, immunization with dry powder formulated peptide patches via skin micropores generated by fractional laser ablation can induce potent CTL responses when combined with a suitable adjuvant.

## Declaration of Competing Interest

RB is head of business development, WB is sales director, and MB is medical scientific director of Pantec Biosolutions AG. RW reports having received grant money from Pantec Biosolutions AG. SS and SD declare that they have no known competing financial interests or personal relationships that could have appeared to influence the work reported in this paper.

## References

[1] Weiss R, Hessenberger M, Kitzmuller S, Bach D, Weinberger EE, Krautgartner WD, et al. Transcutaneous vaccination via laser microporation. J Control Release. 2012;162:391–9.

[2] Terhorst D, Fossum E, Baranska A, Tamoutounour S, Malosse C, Garbani M, et al. Laser-assisted intradermal delivery of adjuvant-free vaccines targeting XCR1+ dendritic cells induces potent antitumoral responses. J Immunol. 2015;194:5895–902.

[3] Chen X, Kim P, Farinelli B, Doukas A, Yun SH, Gelfand JA, et al. A novel laser vaccine adjuvant increases the motility of antigen presenting cells. PLoS One. 2010;5:e13776.

[4] Wang J, Shah D, Chen X, Anderson RR, Wu MX. A micro-sterile inflammation array as an adjuvant for influenza vaccines. Nat Commun. 2014;5:4447.

[5] Chen X, Kositratna G, Zhou C, Manstein D, Wu MX. Micro-fractional epidermal powder delivery for improved skin vaccination. J Control Release. 2014;192:310 –6.

[6] Constantino J, Gomes C, Falcao A, Neves BM, Cruz MT. Dendritic cell-based immunotherapy: a basic review and recent advances. Immunol Res. 2017;65:798 –810.

[7] Chen X, Zeng Q, Wu MX. Improved efficacy of dendritic cell-based immunotherapy by cutaneous laser illumination. Clin Cancer Res. 2012;18:2240 –9.

[8] Scheiblhofer S, Strobl A, Hoepflinger V, Thalhamer T, Steiner M, Thalhamer J, et al. Skin vaccination via fractional infrared laser ablation - Optimization of laser-parameters and adjuvantation. Vaccine. 2017;35:1802 –9.

[9] Stoecklinger A, Grieshuber I, Scheiblhofer S, Weiss R, Ritter U, Kissenpfennig A, et al. Epidermal langerhans cells are dispensable for humoral and cell-mediated immunity elicited by gene gun immunization. J Immunol. 2007;179:886 –93.

[10] Li H, Fan L, Zhu S, Shin MK, Lu F, Qu J, et al. Epilation induces hair and skin pigmentation through an EDN3/EDNRB-dependent regenerative response of melanocyte stem cells. Sci Rep. 2017;7:7272.

[11] Sparwasser T, Hultner L, Koch ES, Luz A, Lipford GB, Wagner H. Immunostimulatory CpG-oligodeoxynucleotides cause extramedullary murine hemopoiesis. J Immunol. 1999;162:2368 –74.

[12] Serre K, Giraudo L, Siret C, Leserman L, Machy P. CD4 T cell help is required for primary CD8 T cell responses to vesicular antigen delivered to dendritic cells in vivo. Eur J Immunol. 2006;36:1386–97.

[13] Ahrends T, Busselaar J, Severson TM, Babala N, de Vries E, Bovens A, et al. CD4(+) T cell help creates memory CD8(+) T cells with innate and help-independent recall capacities. Nat Commun. 2019;10:5531.

[14] Wu CY, Monie A, Pang X, Hung CF, Wu TC. Improving therapeutic HPV peptide-based vaccine potency by enhancing CD4+ T help and dendritic cell activation. J Biomed Sci. 2010;17:88.

[15] Alexander J, Fikes J, Hoffman S, Franke E, Sacci J, Appella E, et al. The optimization of helper T lymphocyte (HTL) function in vaccine development. Immunol Res. 1998;18:79–92.

[16] Cheng WK, Wee K, Kollmann TR, Dutz JP. Topical CpG adjuvantation of a protein-based vaccine induces protective immunity to Listeria monocytogenes. Clin Vaccine Immunol. 2014;21:329 –39.

[17] Stein P, Gogoll K, Tenzer S, Schild H, Stevanovic S, Langguth P, et al. Efficacy of imiquimod-based transcutaneous immunization using a nano-dispersed emulsion gel formulation. PLoS One. 2014;9:e102664.

[18] Rechtsteiner G, Warger T, Osterloh P, Schild H, Radsak MP. Cutting edge: priming of CTL by transcutaneous peptide immunization with imiquimod. J Immunol. 2005;174:2476–80.

[19] Demuth PC, Garcia-Beltran WF, Ai-Ling ML, Hammond PT, Irvine DJ. Composite dissolving microneedles for coordinated control of antigen and adjuvant delivery kinetics in transcutaneous vaccination. Adv Funct Mater. 2013;23:161 –72.

[20] Kadowaki N, Ho S, Antonenko S, Malefyt RW, Kastelein RA, Bazan F, et al. Subsets of human dendritic cell precursors express different toll-like receptors and respond to different microbial antigens. J Exp Med. 2001;194:863 –9.

[21] Molenkamp BG, van Leeuwen PA, Meijer S, Sluijter BJ, Wijnands PG, Baars A, et al. Intradermal CpG-B activates both plasmacytoid and myeloid dendritic cells in the sentinel lymph node of melanoma patients. Clin Cancer Res. 2007;13:2961 –9.

[22] Gursel M, Verthelyi D, Klinman DM. CpG oligodeoxynucleotides induce human monocytes to mature into functional dendritic cells. Eur J Immunol. 2002;32:2617 –22.

[23] Chiang CL, Kandalaft LE. In vivo cancer vaccination: Which dendritic cells to target and how? Cancer Treat Rev. 2018;71:88–101.

[24] Adamus T, Kortylewski M. The revival of CpG oligonucleotide-based cancer immunotherapies. Contemp Oncol (Pozn). 2018;22:56–60.

[25] Ghaffari-Nazari H, Tavakkol-Afshari J, Jaafari MR, Tahaghoghi-Hajghorbani S, Masoumi E, Jalali SA. Improving Multi-Epitope Long Peptide Vaccine Potency by Using a Strategy that Enhances CD4+ T Help in BALB/c Mice. PLoS One. 2015;10:e0142563.

